# Real-Time Multi-Angle Projection Imaging of Biological Dynamics

**DOI:** 10.1101/2020.10.29.355917

**Authors:** Bo-Jui Chang, Etai Sapoznik, Theresa Pohlkamp, Tamara S. Terrones, Erik S. Welf, James D. Manton, Philippe Roudot, Kayley Hake, Lachlan Whitehead, Andrew G. York, Kevin M. Dean, Reto Fiolka

**Affiliations:** Department of Cell Biology, University of Texas Southwestern Medical Center, Dallas, TX 75390, USA; Department of Molecular Genetics, UT Southwestern Medical Center, 6000 Harry Hines Blvd., Dallas, Texas 75390, USA; Lyda Hill Department of Bioinformatics, University of Texas Southwestern Medical Center, Dallas, TX 75390, USA; MRC Laboratory of Molecular Biology, Cambridge, CB2 0QH, UK; Calico Life Sciences LLC, South San Francisco, CA, USA; Walter and Eliza Hall Institute of Medical Research, Parkville, Victoria 3052, Australia; Department of Medical Biology, University of Melbourne, Parkville, Victoria, Australia

## Abstract

We introduce a cost-effective and easily implemented scan unit which enables any camera-based microscope to perform projection imaging from diverse viewing angles. We demonstrate this capability on Lattice Light-Sheet and Oblique Plane Microscopy by rapidly delivering projection images with an uncompromised lateral resolution and high optical contrast. By imaging the sample from one or multiple perspectives, our method enables visualization of rapid biological processes, real time stereoscopic imaging as well as three-dimensional particle localization throughout a cellular volume from just two images. Furthermore, because our projection imaging technique provides intuitive three-dimensional renderings in real-time, it improves microscope usability, allows users to more-readily optimize instrument performance and identify biological phenomena of interest on-the-fly, while also reducing data overhead by a factor of >100. We leverage our rapid projection method to image cancer cell morpho-dynamics and calcium signaling in cultured neurons, to perform three-dimensional localization of genetically encoded nanoparticles, as well as to image orthogonal views of an embryonic Zebrafish heart simultaneously.

## Main Text

Quantitative biological imaging requires Nyquist sampling in both space and time. Yet, owing to the limited acquisition rate of modern laser scanning and camera-based microscopes, many biological processes occur too rapidly to be observed three-dimensionally. In part, this is because volumes are usually acquired in the form of a focal stack, i.e., the sequential acquisition of tens to hundreds of two-dimensional focal planes. However, imaging rates could be orders of magnitude faster if information spanning multiple focal planes could be integrated into a single raster scan or camera exposure. Indeed, multiple projection imaging methods exist^1-3^, but they typically deteriorate lateral resolution, require specialized setups and provide only projections along the optical axis. While a projection from a single direction can be sufficient for sparse samples, multiple viewing angles are desirable on denser samples to provide alternative perspectives and to employ stereoscopic or tomographic approaches to reconstruct the full three-dimensional volume. Nonetheless, enabling variable projection angles in turn requires rotation of the sample, which is impractical for many applications.

Here, we introduce a simple scan unit that converts any camera-based microscope into a projection imaging system that is capable of integrating information from diverse viewing angles. Importantly, our method does not require sample rotation, but instead exploits optical shearing to provide variable viewing directions of the sample. Our projection method can be advantageously combined with Lattice Light-sheet Microscopy (LLSM)^4^ and Oblique Plane Microscopy^5^ (OPM), delivering rapid, high-contrast 2D projections from arbitrary directions with uncompromised lateral resolution. Furthermore, LLSM and OPM require computational shearing of the image data for traditional visualization and quantification (**Figure 1A-C**). Instead, our projection method can perform the required shearing optically during the acquisition of a single camera frame, providing much more intuitive and instantaneous 3D renderings of the sample. For practical matters, this turns out to make such a microscope much more user-friendly, as complex samples can be explored in a more intuitive, epi-fluorescence microscopy view and rapid variation of the projection angle can provide 3D perspective.

**Figure 1 -.**
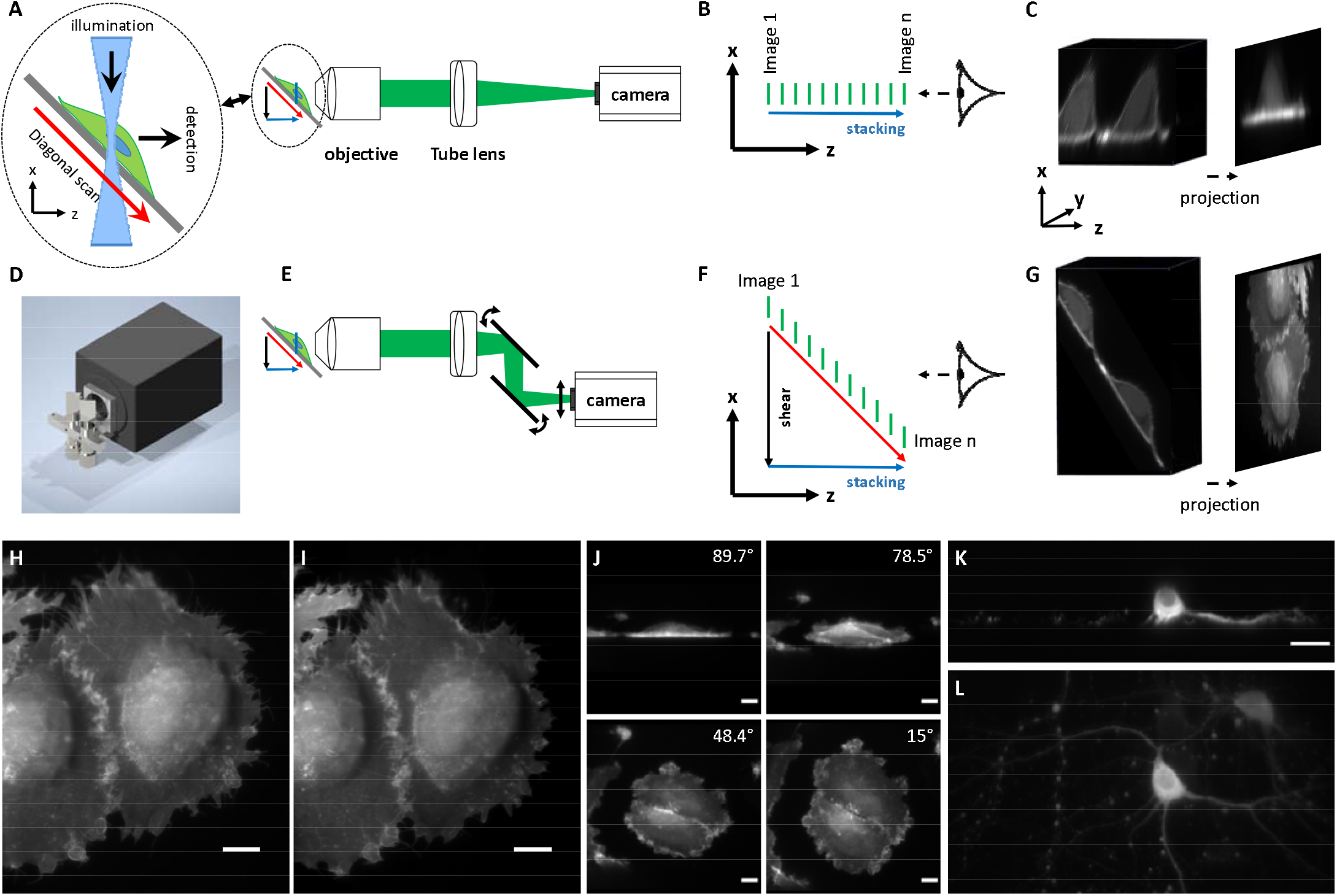
Principle of multi angle projection imaging. **A** In a conventional lattice light-sheet or OPM microscope, the sample is scanned on a diagonal trajectory (red arrow) relative to the detection axis, while being illuminated with a light-sheet (blue line). **B** The images acquired in this way (green tiles) are stacked in the z-dimension one after the other to form a “3D stack”. However, this stack does not reflect the x-component of the scan trajectory (black arrow in A), but only its axial component (blue in **A**); **C** Experimental data stack acquired with LLSM and its projection. **D** CAD rendering of the scan unit attached to the camera. **E** For projection imaging under different viewing angles, a lateral shearing unit consisting of two galvanometric mirrors is added in front of the camera. When the sample is scanned (red arrow), these two mirrors are rotated in synchrony, causing the image to be displaced laterally on the camera (black double headed arrow). F The images acquired this way (green tiles) are stacked in the z (blue arrow, labeled “stacking”) and laterally shifted in the x-direction (black arrow, labeled “shear”). The arrangement of these images corresponds to the natural position of the sections in the sample (see also red arrow). G Same experimental data as in C, but after applying shearing. **C and G** A projection can be computed by numerically summing all the tiles in the 3D stack together or by scanning the sample one or multiple times during one camera exposure. H Projection view of the same MV3 cells expressing AKTPH-GFP, acquired with LLSM-Pro. This view was acquired on a single camera frame. I Numerical projection of a conventional z-stack (334 planes) of MV3 cells labeled with AKT-PH-GFP, acquired with conventional LLSM.J Projection views of another MV3 cell under different shearing parameters (viewing angle measured to the normal of the coverslip), acquired with LLSM-Pro. K single slice view of cultured Neurons, acquired with OPM. L Projection view of cultured neurons, acquired with OPM-Pro. Scale bars: 10 microns.

To implement our projection method, we developed a simple galvanometer-based module that attaches to the microscope camera in a non-perturbative fashion and shears the data optically instead of computationally (**Figure 1 D and Supplementary Figure 1**). Specifically, by sweeping the scan unit synchronously with the acquisition of a z-stack, high-contrast and high-resolution volumetric data can be projected (by taking the sum-intensity) onto a single camera frame (**Figure 1D-G, Supplementary Figure 2, Supplementary Movie 1-2**). Importantly, this allows for a “three-dimensional rendering” of the sample under investigation to be viewed in real-time prior to volumetric “slice-by-slice” imaging (**Supplementary Movie 3**), and provides real-time feedback on instrument alignment and optimization. By changing the magnitude of the scan sweep, projections are obtained from different viewing perspectives in a manner that is mathematically analogous to a shear-warp transform^6^ (**Supplementary Figure 3, Supplementary Movie 4, and Supplementary Note 1**).

As proof of principle, we first imaged mammalian cells using a high-resolution laser scanning OPM^7^ and a sample scanning Field Synthesis variant of LLSM^4, 8, 9^, referred to as OPM-Pro and LLSM-Pro. “Pro” is the shorthand notation for operating the microscope in our projection mode. In both cases, the addition of our scan unit did not cause any significant changes to the optical train, and the microscope could rapidly be switched between volumetric and projection imaging modes. **Figure 1H** shows two MV3 cells labelled with the PI3K biosensor AktPH-GFP and imaged with LLSM-Pro. In contrast, **Figure 1I** shows the same two cells as imaged conventionally with LLSM (a z-stack encompassing 334 slices, computationally sheared, and sum intensity projected). The two images are indistinguishable (**Supplementary Figure 4**), despite the fact that LLSM-Pro acquired the data in a single frame and thus reduced the imaging time and data overhead 334-fold. **Figure 1J** shows a pair of MV3 cells imaged by sweeping the viewing perspective (**Supplementary Movie 4**). To demonstrate the usefulness of our projection method to explore complex samples, we imaged cultured neurons expressing GCaMP6f. **Figure 1K** shows a conventional view as obtained by OPM. While one can identify a cell body and one axon, it is hard to discern the context (i.e. does this neuron have many projections, is it in contact with neighboring neurons), which is not changed when scanning through the sample (**Supplementary Movie 3**). In contrast, imaging with OPM-Pro instantly reveals the entire neuron and its connections in a single view (**Figure 1L**).

To demonstrate the potential for imaging rapid cellular dynamics, we evaluated the formation and retraction of pressure-based blebs in MV3 cells with LLSM-Pro (**Figure 2A-D, Supplementary Movie 5**). Kymographs demonstrate that protrusion dynamics are sampled finely enough to reveal the different phases of both expansion and retraction (**Figure 2B, D**). Here, each bleb has a unique initial protrusion velocity that is followed by a second, slower phase of growth. Interestingly, observation of two different growth phases challenges the canonical model for bleb growth that consists of purely monotonic expansion and retraction phases^10^. In LLSM-Pro, the imaging speed is limited by the rate at which the sample scanned through the light-sheet. In contrast, because 3D scanning in OPM-Pro is galvanometer-based, the imaging speed is limited primarily by the camera frame rate or the available fluorescence signal. Using OPM-Pro, we were thus able to acquire a time series of calcium waves in cultured neurons at video rate (**Figure 2E-F, Supplementary Table 1** and **Supplementary Movie 6-8**). Here, the ability to image volume projections directly allowed us to observe calcium waves propagating through dendritic arbors from different viewing perspectives, which otherwise would have been challenging to acquire with conventional imaging approaches.

**Figure 2 -.**
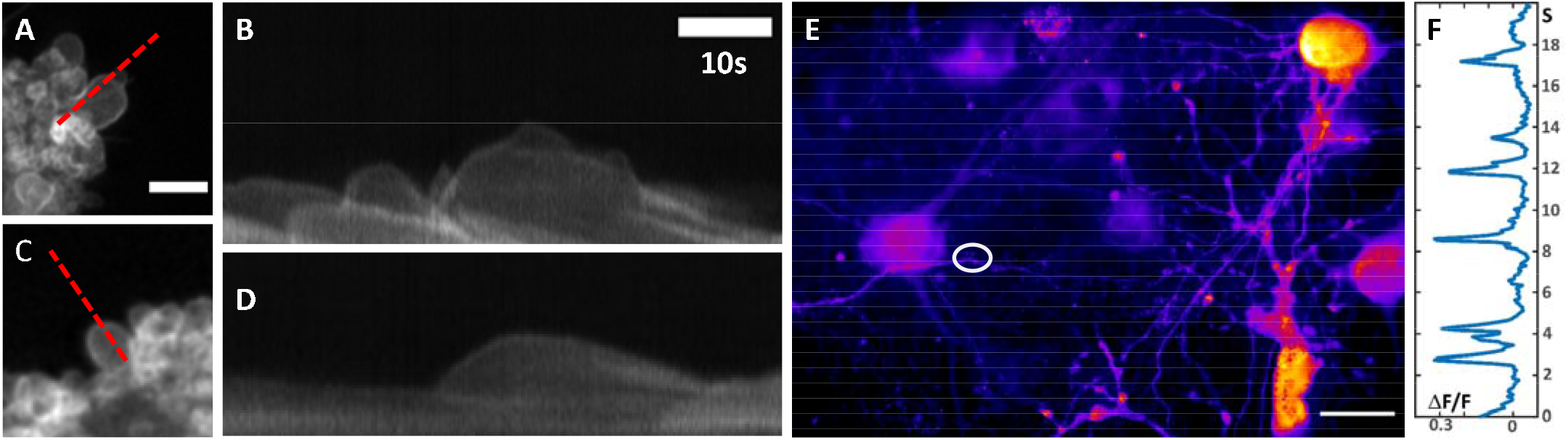
Dynamic projection imaging. **A&C** Blebs on another MV3 cell labeled with AKT-PH-GFP as imaged with LLSM-Pro. **B&D** Kymographs along the dotted lines shown in A&C. Red and magenta lines mark different growth phases in two blebs. E Projection of cultured cortical neurons expressing the calcium indicator GCaMP6f as imaged by OPM-Pro. F Signal trace from the oval region marked in E. Scale bars: A 5 microns; E 20 microns.

The ability to acquire data from multiple perspectives offers several other unique opportunities as well. For example, by rapidly acquiring images from two different viewing perspectives, and overlaying each perspective in a color-coded fashion (here red and cyan), images can be rendered stereoscopically as an anaglyph to the microscope user in real-time (**Figure 3A-B**). Thus, one could operate a microscope equipped with our scan unit in a virtual reality mode where biological specimens are visualized volumetrically (**Supplementary Movie 9**). Additionally, two images acquired from different projection perspectives can encode the information necessary to reconstruct a complete three-dimensional volume. This is most easily demonstrated with sparse objects, which can be localized three-dimensionally via simple triangulation (**Supplementary Note 2-3 and Supplementary Figure 5**). As proof of principle, we localized genetically encoded monomeric nanoparticles^11^ in a fixed MV3 cell from two different views differing by an angle of 18 degrees (**Figure 3C**). Importantly, these inferred locations were in good agreement with a conventional three-dimensional stack that served as a ground truth (**Supplementary Figure 6**). And lastly, for many biological processes, it may be advantageous to obtain two views simultaneously. To achieve this, we added a beam splitter and a second camera that operated without our scan unit and thus acquired a projection along the sample scan direction (**Figure 3D**). This allowed us to simultaneously acquire two orthogonal projections (**Figure 3E**) of the beating heart in a 3 DPF zebrafish embryo at 10 Hz (**Figure 3F**). Due to its rapid motion, acquiring volumetric data of the beating zebrafish heart is challenging. Indeed, acquiring a 3D stack conventionally leads to motion blurring throughout the data, as well as the stripes artifacts when looking from the side (xz or yz view) of the stack (**Supplementary Figure 7**), which either needs to be compensated computationally^12^, via precise synchronization of image acquisition with cardiac dynamics^13^, or by using very rapid acquisition schemes and relatively coarse axial steps sizes^14^. In contrast, our projection method allows us to readily obtain two orthogonal projections with high spatial resolution and no apparent distortions.

**Figure 3 -.**
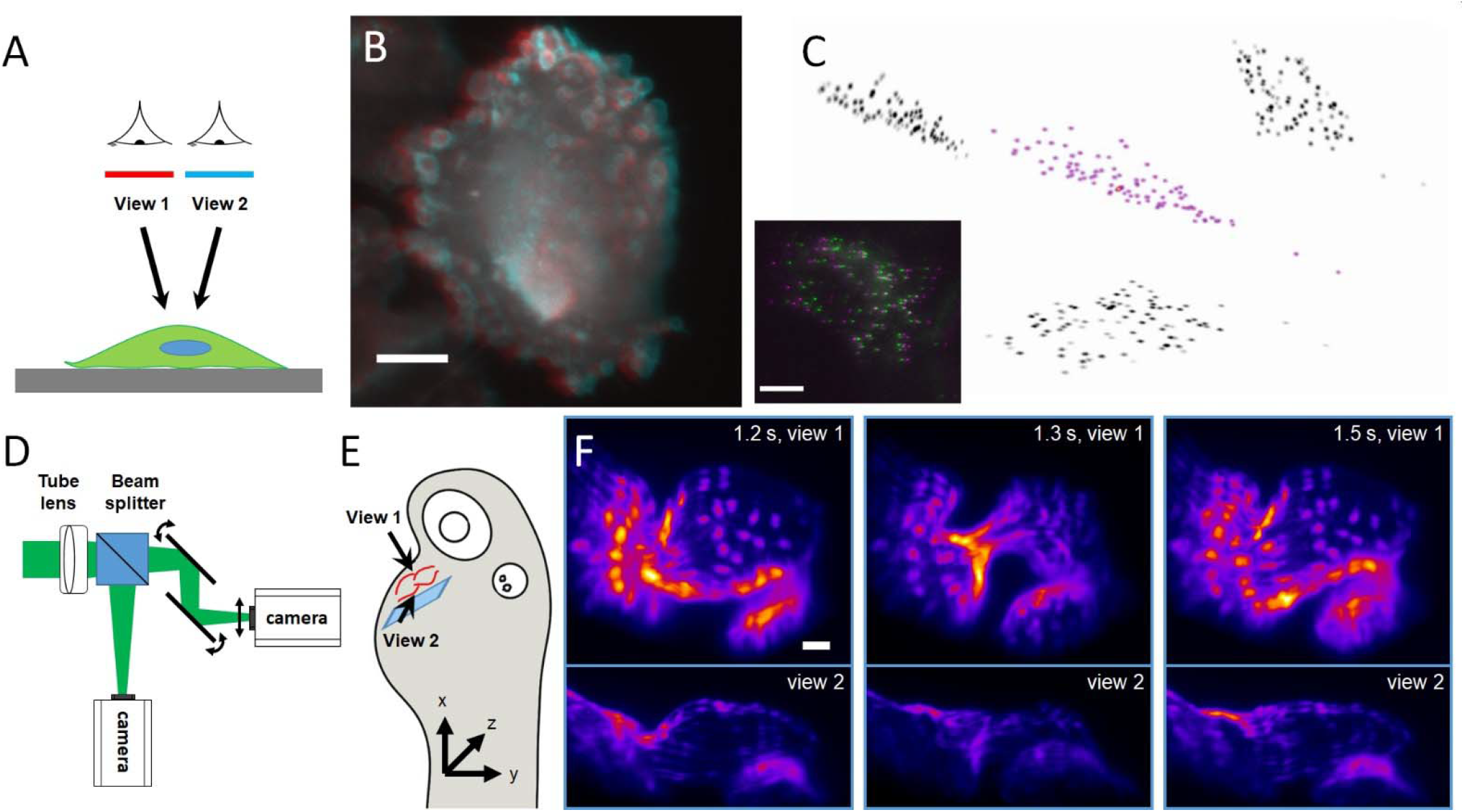
Simultaneous Multi Angle Viewing. **A** Schematic illustration of two viewing directions of a cell on a coverslip. **B** Anaglyph for 3D viewing of a MV3 cancer cells expressing AKT-PH GFP obtained with LLSM-Pro. 3D red cyan glasses are recommended to view this image correctly. The image was created by two different views (15° and −15° to the normal of the coverslip) of the cell at a frame rate of 1Hz. **C** 3D localization from two views of genetically encoded monomeric nanoparticles in a fixed MV3 cell. Magenta shows a rendering of the location in 3D, and maximum intensity projections are shown on the side in black. Small inset shows the two projections (green and magenta) obtained with OPM-Pro. The difference in viewing angle between the two views amounts to 18 degrees. **D** Schematic illustration of a dual camera setup that allows simultaneous acquisition of projections under two different viewing angles. **E** Orthogonal viewing directions under which an embryonic zebrafish heart was imaged. F Three frames of a movie of orthogonal projections of a beating zebrafish heart imaged at a framerate of 10 Hz, obtained with LLSM-Pro. The veins/endothelial tissue of the zebrafish was labeled with GFP (Tg(krdl:GFP)). View 1 is −60 degrees and view 2 is 30 degrees relative to the nominal focal plane of the detection objective. Scale bars: 10 microns in **B-C** and 20 microns in **F**.

These results demonstrate that our scan module can easily augment existing LLSM and OPM imaging systems, and in principle, any camera-based microscope that is equipped with fast z-scanning (**Supplementary Figure 8**). In contrast to previous light-sheet projection^15^, light-field microscopy^16^, and extended depth of focus methods^1, 15, 17^, this is done without any apparent disadvantages as the system is compact, introduces negligible light-losses (<1%), and is fully compatible with normal microscope operation. Of note, for fast projection imaging, sample brightness is an important factor. Here, it is helpful to compare the image formation in our projection method to the acquisition of a traditional z-stack. For fast acquisition times, the virtual exposure time of each individual slice becomes increasingly short (~310 μs in the case of the Zebrafish heart), and this becomes shorter as the volume that is imaged increases. This analogy furthermore illustrates a key difference to approaches that encode axial information into a single PSF; Such PSF engineering approaches are typically limited to small volumes or spread out their PSF over large areas^2^, whereas our projection technique can in principle cover large volumes only limited by the axial scan range while maintaining an invariant PSF. However, our method does so by reducing the fluorescence signal that comes from any emitter when the volume is enlarged.

In conclusion, by exploiting the shear-warp transform, we achieve rotated projections at intermediate angles, which is unlike any previous projection technique. This allowed us to rapidly generate two different perspectives of the sample, which we leveraged for virtual reality viewing of the sample and for the threedimensional localization of sparse objects. Indeed, we foresee applications where our projection method can be used for particle tracking or functional imaging of sparsely labeled neurons in vivo. Because only two images are necessary to reconstruct the imaging volume, this could improve the image acquisition speed by two orders of magnitude compared to conventional mechanisms. With the application of machine learning or tomographic reconstruction, we envision also the reconstruction of more complex, densely labeled specimens. In cases where the axial information can be discarded, our projection method also allows for high-resolution sum projection imaging that is N-fold faster than a traditional light-sheet microscope, where N is the number of Z-slices necessary to acquire a conventional 3D image stack. Despite sacrificing the spatial information in one dimension, such projection imaging methods have proven particularly useful in fields such as neuroscience whereby large volumes must be imaged rapidly in order to resolve the coordinated activity of neuronal circuits. Importantly, because the signal is derived using light-sheet illumination, it inherently has high-contrast and is free from out-of-focus blur. Consequently, we believe that the projection method presented herein will greatly improve research productivity as it transforms relatively non-intuitive yet cutting-edge imaging technologies into user-friendly and interactive data acquisition and hypothesis testing machines.

## Acknowledgments

We would like to thank Dr. David Saucier for the help with the preparation of the zebrafish samples. We would like to thank the Cancer Prevention Research Institute of Texas (RR160057 to R.F.), the National Institutes of Health (T32CA080621 to M.K., F32GM117793 to K.M.D., K25CA204526 to E.S.W., R33CA235254 and R35GM133522 to R.F.) JDM acknowledges support from Fitzwilliam College, Cambridge, through a Research Fellowship.

## Author Contributions

R.F. conceived and build the scan unit. R.F. and B-J.C. designed, built, and operated the microscope. R.F. and A.Y. designed multi-angle research. E.S. and E.S.W. provided MV3 cells. T.P. and T.S.T. provided primary neurons. R.F., J.D.M derived relationship between rotated projections and sheared projections of volumes. L.W. and K.H developed animations. J.D.M. derived equations for localization precision. P.R. performed the 3D reconstruction from two projections. R.F. and K.M.D. wrote the manuscript. All authors read and provided feedback on the final manuscript.

## Competing Interests

R.F. filed a patent for the scan unit and its applications to microscopy. K.M.D. and R.F. have an investment interest in Discovery Imaging Systems, LLC.

## Data Availability

All data presented herein are available from the corresponding author upon request.

## Code Availability

The MATLAB scripts used in this manuscript are available under: https://github.com/AdvancedImagingUTSW

## Methods

### Scan Unit

The scan unit consist of two galvanometric mirrors (6220H, Cambridge technologies), which are mounted in a 3D printed adapter (**Supplementary Figure 2**), which is directly attached to a scientific CMOS camera in both a Field Synthesis microscope and an OPM. Once the shearing unit is attached, the camera has to be shifted by about one inch laterally and axially to image the same focal plane. We found the correct position by using an alignment laser that can be injected into the detection path (see https://github.com/AdvancedImagingUTSW/LaserAlignmentTool). For projection imaging, the sample scanning piezo for LLSM, or the mirror galvanometer that scans the light-sheet and de-scans the fluorescence in an OPM, needs to perform at least one scan, or an integer multiple, during a single camera exposure. To accomplish this, our microscope software generates a sawtooth signal that is synchronized to the camera exposure. This signal then drives the position of the galvanometer-based scan unit, as described in detail below.

### Oblique Plane Microscope

Our OPM was inspired by the work of Bin Yang et al.^5^ and included a novel glass-tipped tertiary^18^ objective that improves the microscope resolution and field of view, as described previously^7^. The only modification to the imaging system was the introduction of the scan can unit into the detection path. The sawtooth signal was applied to the galvanometric mirror (which performs the light-sheet scanning and fluorescence de-scanning), and a scaled version of this signal was sent to the galvo mirrors of the shearing unit. For the latter, we did not use scaling amplifiers, but set up a duplicate signal output on the FPGA card (PCIe-7852, National Instruments) that could be multiplied by a scalar factor. This performs the same functionality as an external scaling amplifier but allowed changes of the shearing factor on the fly. Cells were plated on standard 35 mm diameter #1.5 thickness glass bottomed dishes. Neurons were imaged in eight well plates.

### Lattice Light-Sheet Microscopy

All LLSM and Field Synthesis experiments were performed as previously described, with the exception that the scan unit was introduced into the detection path^9^. The sawtooth signal from the control software was used to control the sample scanning piezo (P621.1CD, Physik Instrumente), and the position signal from the piezo unit was conditioned with a scaling amplifier (SIM983, Stanford Research Systems), and used to control the position of the galvanometer-based scan unit. Such an approach guaranteed that the shearing unit moved synchronously with the sample scan. Cells were plated on 5 mm coverslips and mounted in a custom sample holder for imaging^6^. Zebrafish embryo was embedded in 1% agarose in a FEP (Fluorinated Ethylene propylene) tube (FEP HS .029 EXP/ .018 REC, Zeus), and then mounted in a custom sample holder. For both types of samples, coverslip and FEP tubes, the scanning direction was inclined by 30 degrees to the nominal focal plane of the detection objective.

### Mammalian Cell Culture

Metastatic melanoma (MV3, a kind gift from Dr. Peter Friedl, MD Anderson Cancer Center) cells were cultured in DMEM supplemented with 10% fetal bovine serum and penicillin/streptomycin and maintained at 37 degrees Celsius with 5% CO_2_ atmosphere. MV3 cells were transduced using the pLVX-IRES-PURO and pLVX-IRES-NEO lentiviral systems.

### Primary Neurons

Animal Care Rat primary neurons were obtained in accordance to protocols approved by the University of Texas Southwestern Medical Center Institutional Animal Care and Use Committee (IACUC). Cell Culture and Labeling Male and female embryonic day 18 (E18) primary cortical neurons were prepared from timed pregnant Sprague Dawley rats (Charles River Laboratories, Wilmington, MA) as previously described^19^. Embryonic cortices were harvested, neurons dissociated, and plated on 35 mm glass bottom culture dishes (MatTek P35G-1.5-10-C)), coated with poly-D-lysine, at a density of 0.8 million neurons per dish. Neurons were cultured in completed Neurobasal medium (Gibco 21103049) supplemented with 2% B27 (Gibco 17504044), 1 mM glutamine (Gibco 25030081), and penicillin streptomycin (Gibco 15140148) at 37°C in a 5% CO2 environment. At day in vitro (DIV) 2 neurons were treated with AraC to prevent overgrowth of glia cells. Half of the culture media was renewed twice a week. On DIV5 neurons were infected with lentivirus encoding GCaMP6f. The lentivirus was generated by co-transfecting HEK 293 T cells with psPAX2, pMD2G (kindly provided by Didier Trono, Addgene numbers 12260, 12259), and pLV-GCaMP6f^20^. Neurons were utilized for live-imaging at DIV12-16.

### Data Analysis

Bleb kympgraphs were analyzed with ImageJ. The Neuron firing trace was extracted in Matlab. Most data visualization and analysis was performed in ImageJ.

### Deconvolution and Post-Processing

Projection data under different angles was compressed in one direction using the imresize function in Matlab. For 3D localization, particles were detected using U-track and the z-location was computed as described in **Supplementary Note 3** using a Matlab script. For 3D and 2D deconvolution, the Matlab Blind deconvolution routine was used with an experimentally measured point spread function. In the 2D deconvolution of the projection data, temporal averaging was performed in some cases as indicated in Supplementary Table 1 to suppress some spurious noise patterns that arose from deconvolution. For all temporal analysis of data (kymographs, firing traces) non deconvolved and non averaged data was used.

### Animal Specimens

All animal protocols were approved by local Institutional Animal Care and Use Committees (IACUC) as directed by the National Institutes of Health, and strictly followed. This included APN 2016-101805 (to Dr. Stephen Skapek, UT Southwestern Medical Center).

