## Supplementary Information for "Real-Time Multi-Angle Projection Imaging of Biological Dynamics"

#### Affiliation

**Supplementary Notes**

**Supplementary Note 1: Equivalence between the projection from a rotated and a sheared volume.**

For the application in LLSM and OPM, our shear unit can perform the necessary shearing to exactly undo the lateral displacement that occurs during the scanning process in those two microscopes. As such, the resulting projection is expected to be equivalent to the numerically sheared and projected 3D data coming from such a microscope. But what happens if the shear unit performs a larger or smaller shearing angle than what is necessitated by the microscope? This general case, i.e shearing by an arbitrary angle and subsequent projection, is investigated below. Consider, without loss of generality, shearing along the x-axis by a and rotating in a positive sense around the z-axis by Θ. The rotation matrix is given by

$$R = \begin{bmatrix} \cos\theta & -\sin\theta \\ \sin\theta & \cos\theta \end{bmatrix}$$

and the shear matrix by

$$S = \begin{bmatrix} 1 & \alpha \\ 0 & 1 \end{bmatrix}$$

In both cases, we wish to project along the y-axis, for which the projection matrix is

$$P = \begin{bmatrix} 1 & 0 \\ 0 & 0 \end{bmatrix}$$

Consider now the projection of the rotation matrix,

$$PR = \begin{bmatrix} 1 & 0 \\ 0 & 0 \end{bmatrix} \begin{bmatrix} \cos\theta & -\sin\theta \\ \sin\theta & \cos\theta \end{bmatrix} = \begin{bmatrix} \cos\theta & -\sin\theta \\ 0 & 0 \end{bmatrix}$$

And the projection shear matrix,

$$PS = \begin{bmatrix} 1 & 0 \\ 0 & 0 \end{bmatrix} \begin{bmatrix} 1 & \alpha \\ 0 & 1 \end{bmatrix} = \begin{bmatrix} 1 & \alpha \\ 0 & 0 \end{bmatrix}$$

If we multiply PS by $\cos\theta$, we can achieve equality if $\alpha = -\sin\theta / \cos\theta = -\tan\theta$.

The multiplication of $\cos\theta$, has the effect of scaling the output:

$$MPS = \begin{bmatrix} Sx & 0 \\ 0 & Sy \end{bmatrix} \begin{bmatrix} 1 & 0 \\ 0 & 0 \end{bmatrix} \begin{bmatrix} 1 & \alpha \\ 0 & 1 \end{bmatrix} = \begin{bmatrix} Sx & \alpha Sx \\ 0 & 0 \end{bmatrix}$$

With $\alpha = -\tan\theta$ and Sx=$\cos\theta$, and M is the scaling matrix. However, in practice, the shearing unit only performs the shearing, but not the scaling operation, i.e.

$$PS = \begin{bmatrix} 1 & 0 \\ 0 & 0 \end{bmatrix} \begin{bmatrix} 1 & \alpha \\ 0 & 1 \end{bmatrix} = \begin{bmatrix} 1 & \alpha \\ 0 & 0 \end{bmatrix}$$

This has the following implications: Applied to lattice and OPM data, if we just undo the amount of shearing imparted by the diagonal scan trajectory, the projection is equivalent to a computationally sheared and projected dataset. However, as we go away from the ideal shear parameter (such that the projected view corresponds to the projection of a properly sheared LLSM or OPM data volume), the resulting projection image will be stretched in one dimension by $1/\cos\theta$, compared to a projection obtained from a volume that is numerically rotated. As such, the scaling operation (i.e. a squeezing of the image in one dimension) has to be done computationally, to make the projected views equivalent. For viewing purposes and for small angles (i.e. approximating $\cos\theta \sim 1$ ), however, this might be ignored.

**Supplementary Note 2: Precision of particle localization using projected views.**

Obtaining two views under different viewing angles allows triangulation of the 3D location of objects, provided the same object can be localized in each view. If this is the case, a simple reconstruction algorithm is schematically shown in Supplementary Figure 5. The projection of two sheared volumes are taken, which have a shear angle of $\theta$ and - $\theta$ relative to the true, unsheared volume. One can find that the height of particles in this volume is given by: $h = \Delta x/2\tan(\theta)$. Δx is the apparent displacement of an object between the two projections.

We wish to calculate the theoretical 3D localization precision for a particle, given two views acquired using a our projection method. We assume that the PSF is well-approximated by an ellipsoidal Gaussian function and that there is no background. Under these assumptions, the Thompson–Mortensen localization variance in each dimension is given by

$$\sigma^2 = \frac{16}{9}\frac{s^2 + a^2/12}{N},$$

where $s$ is the standard deviation of the Gaussian function approximating the PSF, $a$ is the pixel width and $N$ is the number of photons collected[21]. The localization precision, $\sigma$, is given by the positive square root of this quantity.

For the field synthesis light sheet microscope used in this work, PSF measurements on sub-diffraction-sized beads provide full-widths at half-maximum of approximately in $x$, in $y$ and in $z$, corresponding to $s_x$ = 123 nm, $s_y$ = 123 nm and $s_z$ = 318nm. All acquisitions have a pixel size of 104 nm. In a projection acquisition, the PSF is rotated by an angle, $\theta$, around the $x$-axis and also stretched in the $y$ direction by a factor of $1/\cos\theta$. Hence, in the projection view (Pro shorthand for our projection method):

$$s_{x'}^{\mathrm{Pro}} = s_x,$$
$$s_{y'}^{\mathrm{Pro}} = \frac{1}{\cos\theta}\sqrt{\frac{s_z^2 s_y^2}{\left(s_z^2\cos^2\theta + s_y^2\sin^2\theta\right)}},$$

where $x'$ and $y'$ refer to the horizontal and vertical position in the camera frame, not 3D space.

Given two estimators, $\widehat{\alpha}_1$ and $\widehat{\alpha}_2$, of $\alpha$, with variances $\sigma_1^2$ and $\sigma_2^2$, then, for the combined estimator

$$\widehat{\alpha} = \frac{\sigma_2^2\,\widehat{\alpha}_1 + \sigma_1^2\,\widehat{\alpha}_2}{\sigma_1^2 + \sigma_2^2},$$

the variance of $\widehat{\alpha}$ is given by

$$\frac{\sigma_1^2\sigma_2^2}{\sigma_1^2 + \sigma_2^2}.$$

For simplicity, we consider our projection acquisition scheme where both views are rotated by $\pm\theta$ from the conventional deskewed view. We further assume that both views correspond to the same number of integrated photons. Hence, per dimension, the localization precision in each view is equal, e.g. $\sigma_{x'_1}^{\mathrm{Pro}} = \sigma_{x'_2}^{\mathrm{Pro}}$.

As the apparent $x$ co-ordinate is not changed by our projection acquisition procedure, the true $x$ value can be estimated by pooling both $x'$ measurements. Similarly, the true $y$ value is estimated by considering the mean of the two $y'$ measurements. The $z$ location of the particle is given by

$$z = \frac{y'_2 - y'_1}{2\tan\theta}.$$

Hence,

$$\begin{aligned}
\sigma_x^2 &= \frac{\left(\sigma_{y'}^{\mathrm{Pro}}\right)^2}{2} \\
\sigma_y^2 &= \frac{\left(\sigma_{y'}^{\mathrm{Pro}}\right)^2}{2} \\
\sigma_z^2 &= \frac{\left(\sigma_{y'}^{\mathrm{Pro}}\right)^2}{2\tan\theta}.
\end{aligned}$$

We can compare this with a hypothetical situation in which two views at an angle $\pm\theta$ from the conventional deskewed view are acquired through a proper rotation of the volume, rather than a projection acquisition. In this situation, the stretching by $1/\cos\theta$ does not occur, giving

$$\begin{aligned}
s_{x'}^{\mathrm{rot}} &= s_x, \\
s_{y'}^{\mathrm{rot}} &= \sqrt{\frac{s_z^2 s_y^2}{\left(s_z^2\cos^2\theta + s_y^2\sin^2\theta\right)}}.
\end{aligned}$$

Here, the $x$ value is estimated as before and so has the same variance, but now

$$\begin{aligned}
\sigma_y^2 &= \frac{\left(\sigma_{y'}^{\mathrm{rot}}\right)^2}{2\cos\theta}, \\
\sigma_z^2 &= \frac{\left(\sigma_{y'}^{\mathrm{rot}}\right)^2}{2\sin\theta}.
\end{aligned}$$

Supplementary Figure 9 shows localization precisions for our projection method and rotated views over the angular range ±90°, given 500 photons per view. As expected, at small stereo angles the localization precisions are approximately equal for our projection method and rotated views, as at small angles $\cos\theta \sim 1$ and $\sin\theta \sim \tan\theta$. Intuitively, this can be understood as a result of the difference between our projection acquisition and rotated acquisition being the stretching by $1/\cos\theta$, which is $\sim 1$ for small angles. We would also expect both the $y$ and $z$ localisation precisions in the projection case to tend towards infinitely poor at ±90°, as here the image is stretched infinitely far, which is indeed observed. While the rotated case is always better than the our projection case, the difference in precision over 'usable' angles (i.e. ±45°) is only of the order of a few nanometers. Finally, we note that the stereo angle at which equal localization precision in $x$ and $y$ is achieved is ±45° in both our projection and rotated cases, irrespective of photon number ($N$) or inherent PSF widths ($s$). However, a ±45° stereo view is not necessarily the best approach if the goal is to instead minimize the total localization precision

$$\sigma = \sqrt{\sigma_x^2 + \sigma_y^2 + \sigma_z^2},$$

which does depend on the inherent PSF widths.

#### Supplementary Note 3: particle localization using projected views.

Diffraction limited objects where detected on the original projection and projection of the sheared volume using previously published approach[22]. The vertical shift induced by the volume shearing was computed through the vertical association of detected object across projections through the resolution of a linear assignment problem expressed as:

$$\underset{\{a_{ij}\}}{argmin} \sum_{i\in\Omega, j\in\Omega'} c_{ij}a_{ij} \;\; s.t. \sum_{i\in\Omega} a_{ij} = 1 \; and \sum_{j\in\Omega'} a_{ij} = 1,$$

where $a_{ij} \in \{0,1\}$ denotes the assignment of the ith detection in the original projection to the jth measurement in the projection of the sheared volume, and $c_{ij}$ is defined as the vertical distance as

$$c_{ij} = \begin{cases} abs(\boldsymbol{y}_i - \boldsymbol{y}_j'), & \boldsymbol{x}_i < r_x \\ +\infty, & x \geq r_x \end{cases}$$

Where $\boldsymbol{y'}_j$ denotes the position in the projection of the sheared volume and $r_x$ is a lateral gating parameter set to 3. For each optimal pair, the associated depth $z_i$ is estimated as $z_i = c_{ij}/\tan(\alpha)$ where alpha is the shearing angle.

The ground truth was computed using the same detection approaches implemented in 3D and aligned using a gradient based optimization. An estimated position is considered as a true positive if it is matched to an object in the ground truth with distance smaller than 575nm. The associated Jaccard Index (computed as TP/(|Ω|+|Ω'|-TP) where TP is the count of true positives) is 0.56.

**Supplementary Figures**

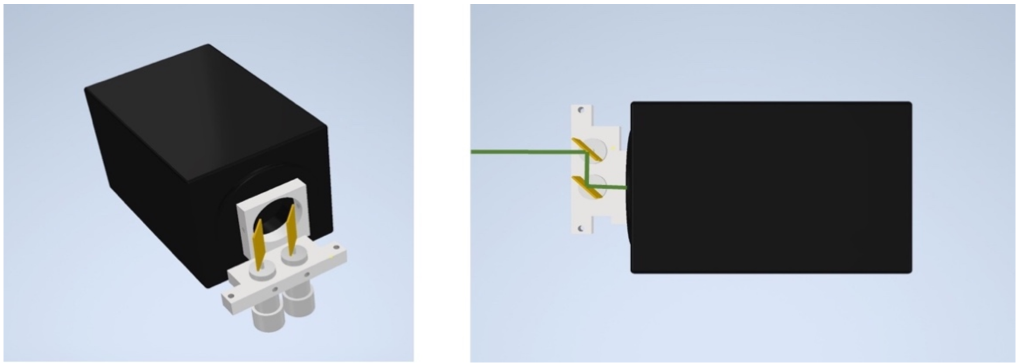

**Supplementary Figure 1 – Design of the Scan Unit.** CAD rendering of the galvanometric scan unit (gray), which attaches to the c-mount of a sCMOS camera (black). Green line shows the path of the fluorescence light over the two galvo mirrors (gold).

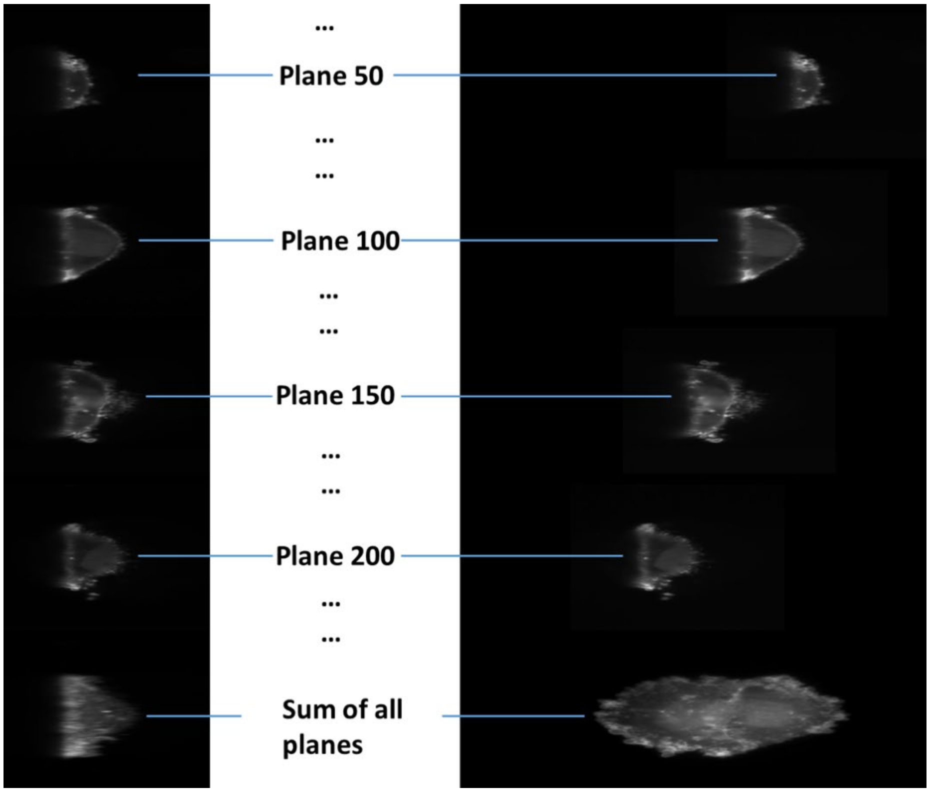


**Supplementary Figure 2 - Illustration of projection-based imaging.** Left shows a conventional stack (every 50th plane shown), as it would be acquired by a lattice light-sheet or oblique plane microscope. Bottom shows the summation of all slices. On the right side, a stack is shown where the appropriate amount of lateral image displacement was added with our shearing unit (every 50th plane is shown). Bottom shows the summation of all slices.

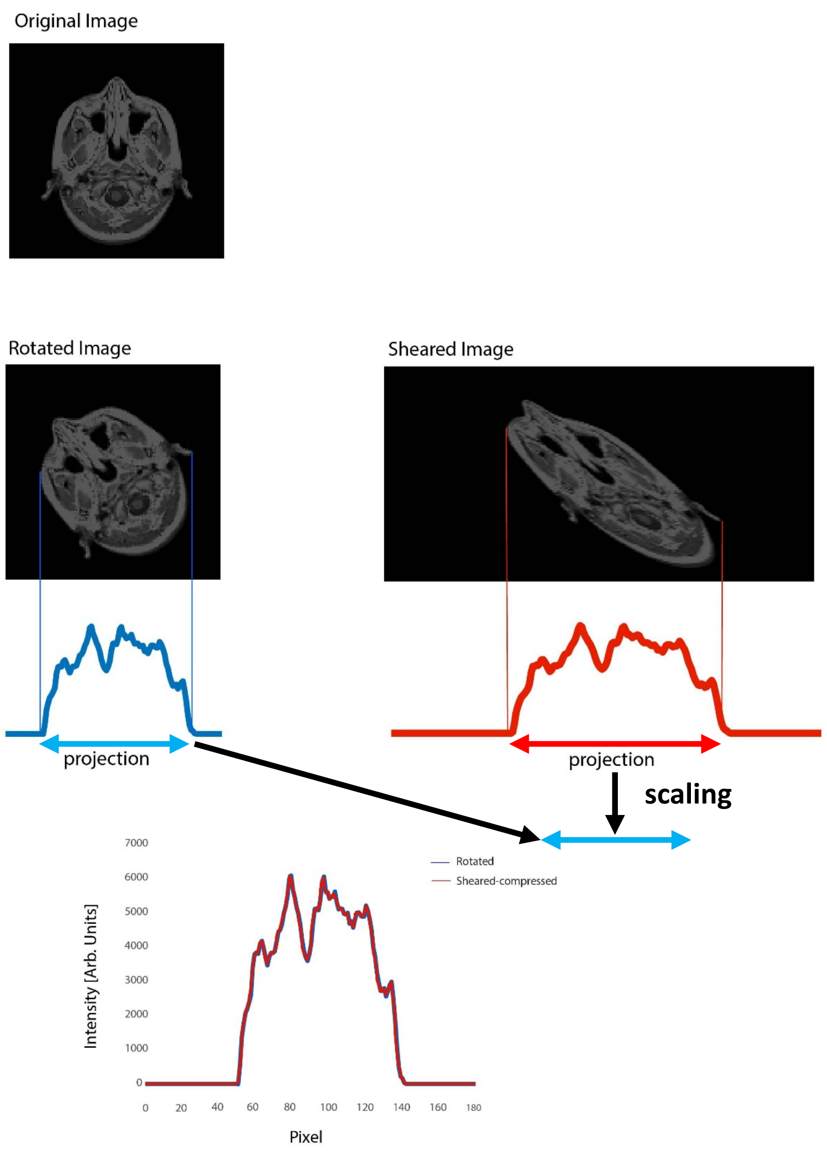


**Supplementary Figure 3 - Equivalence of projection from a rotated and a sheared volume.** Numerical simulation of the rotation of a 2D image and subsequent sum-projection, and comparison to numerical shearing of a 2D image and subsequent sum-projection. To compare the two line-profiles, the projection from the sheared image was compressed to the same length as the projection from the rotated image. Small differences between the two profiles may have arisen from numerical errors during the shearing and rotation operations.

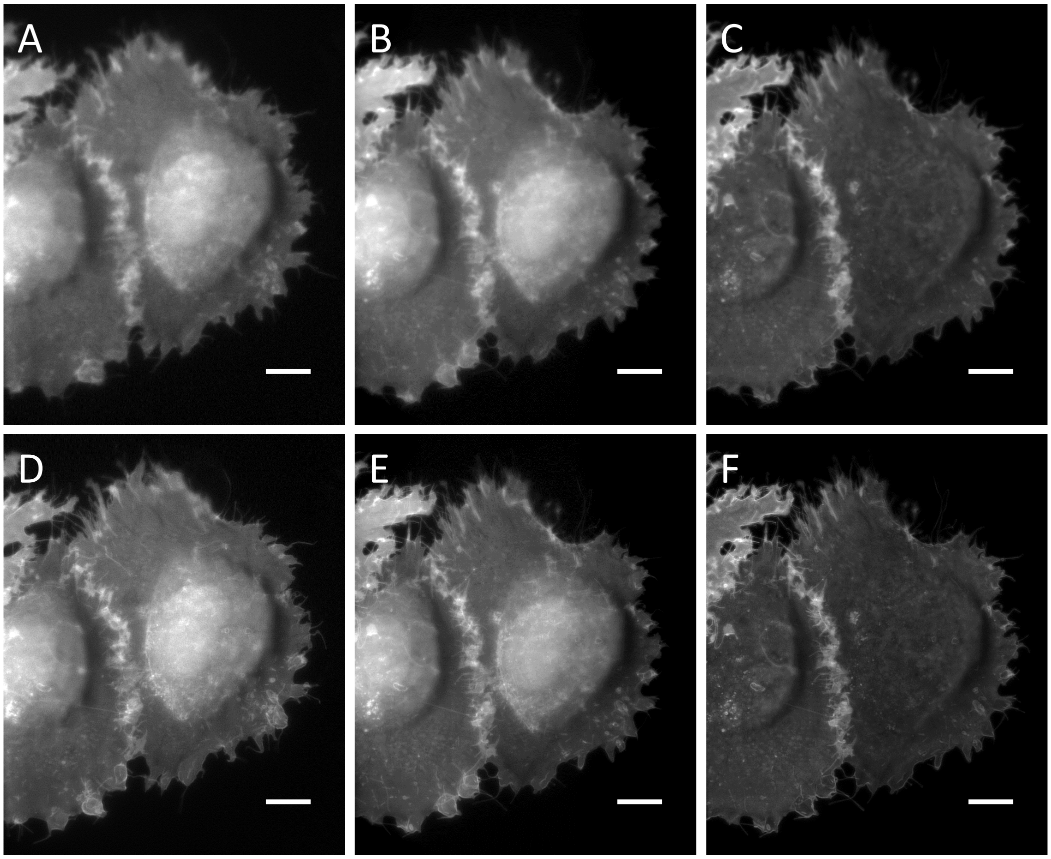


**Supplementary Figure 4 - Comparison of our projection method and numerical projections. A** Single projection obtained with LLSM-Pro. **B** Sum projection of a conventional 3D stack. **C** Maximum intensity projection of a conventional 3D stack. **D** 2D deconvolution of a projection obtained with LLSM-Pro. **E** Sum projection of conventional 3D stack after 3D deconvolution. **F** Maximum intensity projection of conventional 3D stack after 3D deconvolution. Scale bars: 10 microns.

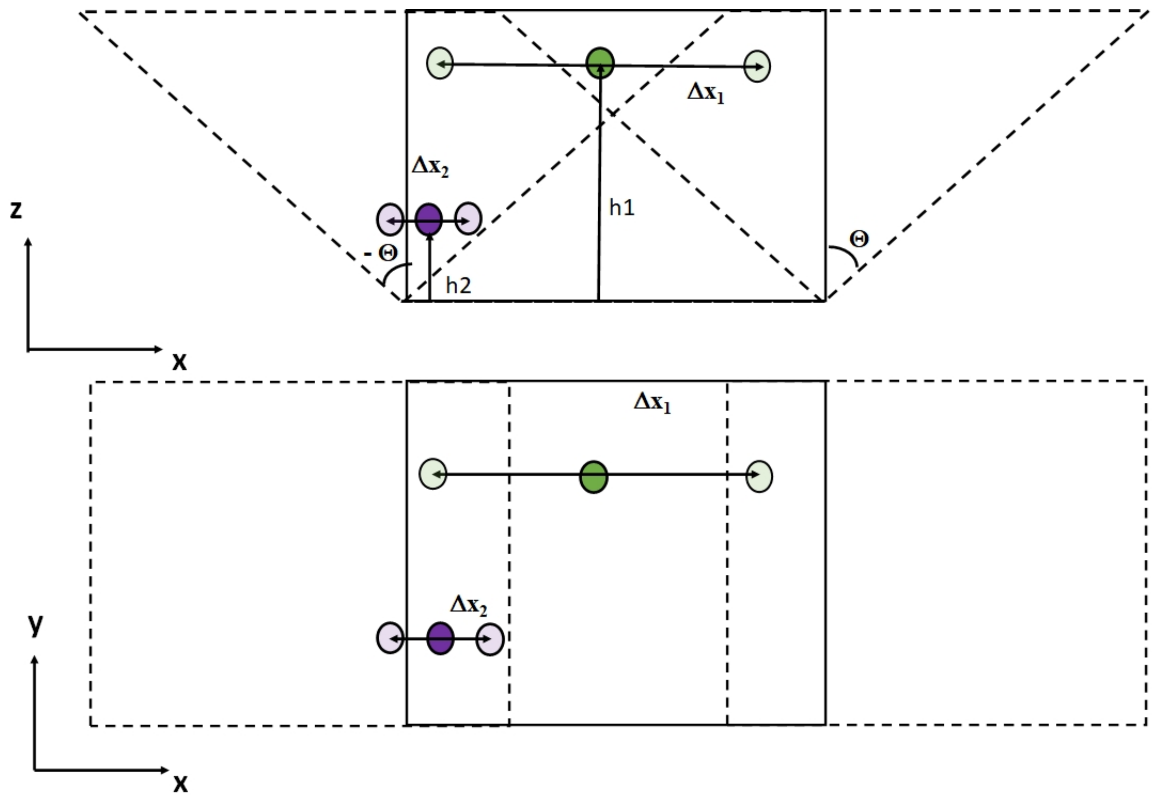


**Supplementary Figure 5 - Determination of the z-position from two projections. Top:** Two volumes sheared by $\pm\theta$ (dotted lines) and an unsheared volume (solid lines) are shown. In these three volumes, two particles, green and magenta, located at different heights are shown. Solid color depicts the particle's true position in the unsheared volume, and transparent colors depicts the particle's position in the sheared volumes. **Bottom:** Projection of the two volumes. The particle's displacement in the x direction, Δx, is proportional to their height.

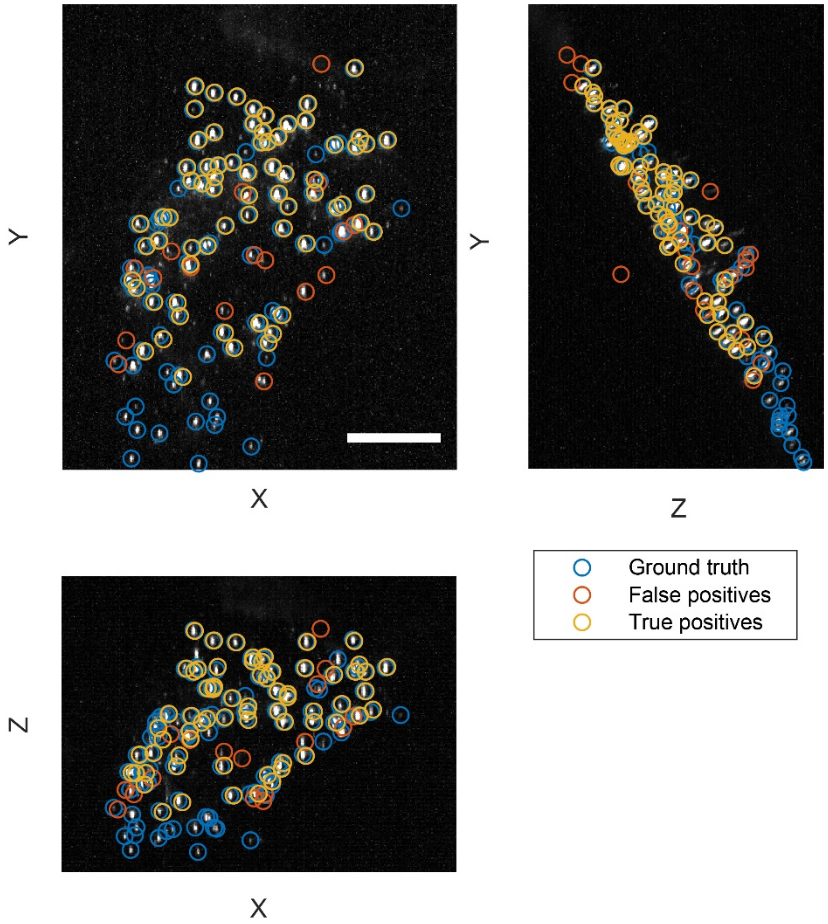


**Supplementary Figure 6 - 3D localization from two views compared with ground truth data from a conventional 3D stack.** Maximum intensity projection of a ground truth data set of genetically encoded monomeric nanoparticles in a fixed MV3 cell (gray) and the localizations of these particles in the 3D stack (blue circles). Yellow circles mark where the reconstruction from two projections overlapped, whereas red circle mark reconstructions that do not match up with the ground truth. Scale bar: 10 microns.

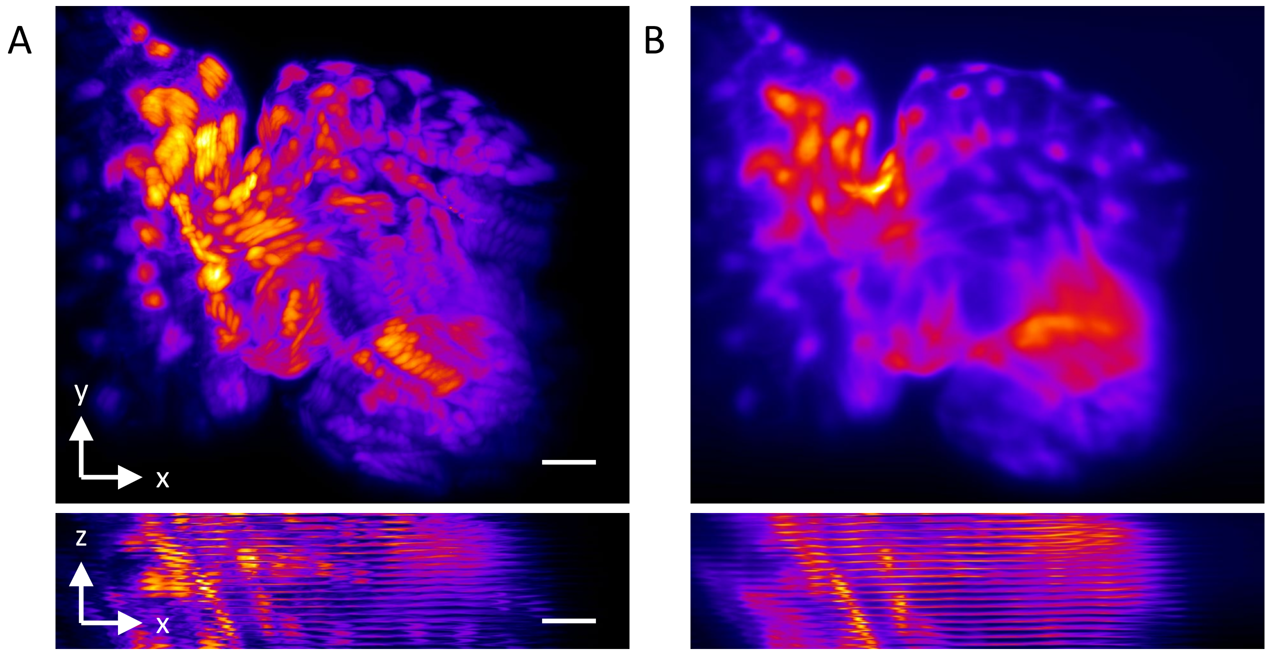


**Supplementary Figure 7 - Zebrafish heart imaging by acquiring conventional 3D stack.** The same zebrafish heart shown in Figure 3F was imaged with a conventional 3D z-stacking. A maximum intensity projection (**A**) and a sum projection (**B**) of the stack in xy (top row) and xz (bottom row) are shown. The exposure time is 25ms (40Hz frame rate), the z step size is 0.312 microns, and there are 322 z slices in the stack. Scale bars: 20 microns.

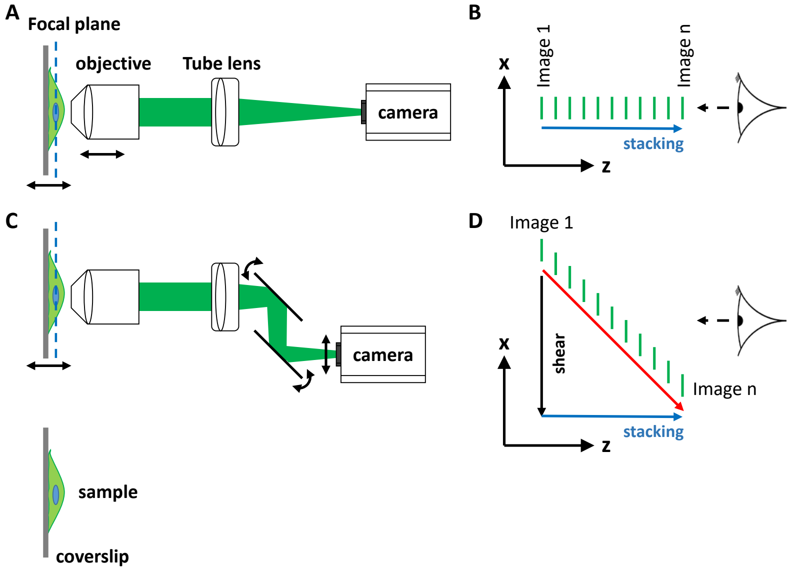


**Supplementary Figure 8 - Working principle of projection imaging in a conventional epifluorescence microscope. A** in a conventional fluorescence microscope, the sample (cell, green translucent) *or* the objective is scanned rapidly to form a 3D stack (black arrows) **B** The images acquired in this way (green tiles) are stacked in the z-dimension one after the other to form a "3D stack". If this dataset is viewed as a projection (dashed arrow), projection of the sample along the optical axis (z) will appear. **C** in our method, a lateral shearing unit consisting of two galvanometric mirrors is added in front of the camera. When the sample is scanned, these two mirrors are rotated in synchrony, causing the image to be displaced laterally on the camera (black double headed arrow). **D** The images acquired this way (green tiles) are stacked in the z (blue arrow, labeled "stacking") and laterally shifted in the x-direction (black arrow, labeled "shear"). In either case (**B** or **D**), a projection can be computed by numerically summing all the tiles in the 3D stack together (forming a "sum projection"). Such a summation can alternatively be formed by scanning the sample one or multiple times during one camera exposure. In this scenario, the sum of all the tiles shown in **B** or **D** will directly appear on the camera as a projection. In our case with the additional shear, the resulting projection appears from a different viewing angle, allowing the user to see different instantaneous views of the sample.

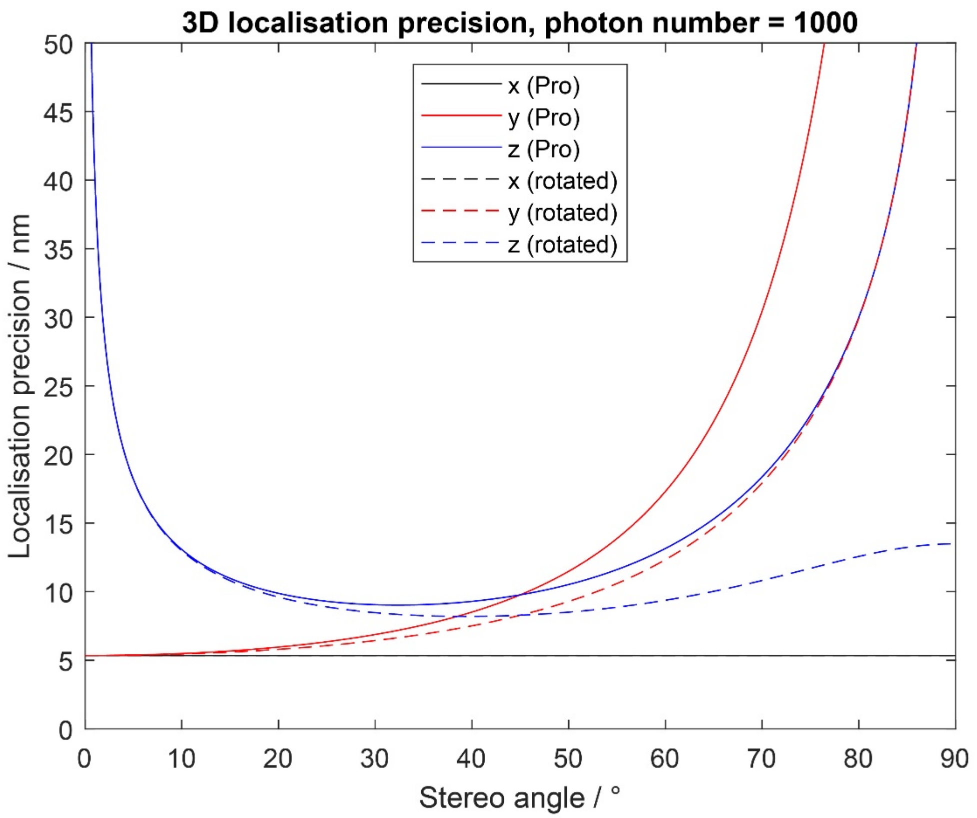


**Supplementary Figure 9 - Thompson–Mortensen localization precisions for two views obtained by our projection method or rotated view imaging.** Each view is ± the stereo angle from the conventional deskewed view, calculated using the bead FWHM measurements described in the main text. As there is no change along the $x$ direction in either method, the $x$ localisation precision is equal in all situations. In both cases, the stereo angle at which equal $y$ and $z$ precision is achieved is ±45°.

### Supplementary Tables

| Figure | microscope | Exposure time | Acquisition rate | Voxel size | Laser power | Post processing |
|---|---|---|---|---|---|---|
| 1C&G | Field Synthesis | 25ms | NAN | 104x104x 300 $nm^3$ | 80µW | Raw data |
| 1H | Field Synthesis | 25ms | NAN | 104x104x 300 $nm^3$ | 80µW | 2D deconvolution |
| 1I | Field Synthesis | 90ms | 5Hz | 104x104$nm^2$ | 80µW | 2D deconvolution, two frame average |
| 1J | Field Synthesis | 90ms | 5Hz | 104x104$nm^2$ | 80µW | Raw data |
| 1K-L | OPM | 100ms | 10Hz | 115x115$nm^2$ | 410 µW | Raw data |
| 2A-D | Field Synthesis | 90ms | 5Hz | 104x104$nm^2$ | 80µW | 2D deconvolution (A,C) and raw data (B,D) |
| 2E-F | OPM | 40ms | 12.5Hz | 115x115$nm^2$ | 750µW | Raw data |
| 3B | Field Synthesis | 250ms | 1Hz | 104x104$nm^2$ | 50µW | anaglyph |
| 3C | OPM | 400ms | NA | 115x115$nm^2$ | 410µW | Raw data and 3D reconstruction |
| 3F | Field Synthesis | 100ms | 10Hz | 104x104$nm^2$ | 30µW | Raw data |
| Video 1 | Animation | NA | NA | NA | NA | NA |
| Video 2 | Field Synthesis | 100ms | 10Hz | 104x104$nm^2$ | 80µW | Raw data |
| Video 3 | OPM | 100ms | 10Hz | 115x115$nm^2$ | 410µW | Raw data |
| Video 4 | Field Synthesis | 90ms | 5Hz | 104x104$nm^2$ | 80µW | 2D deconvolution and Gaussian filter |
| Video 5 | Field Synthesis | 90ms | 5Hz | 104x104$nm^2$ | 80µW | 2D deconvolution |
| Video 6 | OPM | 40ms | 12.5Hz | 115x115nm | 750µW | 2D deconvolution |
| Video 7 | OPM | 50ms | 20Hz | 115x115$nm^2$ | 750µW | Gaussian filter (sigma 1 pixel) |
| Video 8 | OPM | 50ms | 20Hz | 115x115$nm^2$ | 1.65mW | 2D deconvolution and Gaussian filter |
| Video 9 | Field Synthesis | 250ms | 1Hz | 104x104$nm^2$ | 50µW | Raw data |

| | | | | | | |
|---|---|---|---|---|---|---|
| **Video 10** | **Field Synthesis** | **100ms** | **10Hz** | **104x104nm$^2$** | **30µW** | **Raw data** |

**Supplementary Table 1 - Image acquisition parameter and post-processing of the image data.** When a Gaussian filter was used, a width (sigma) of one pixel was applied.

**Supplementary Movies**

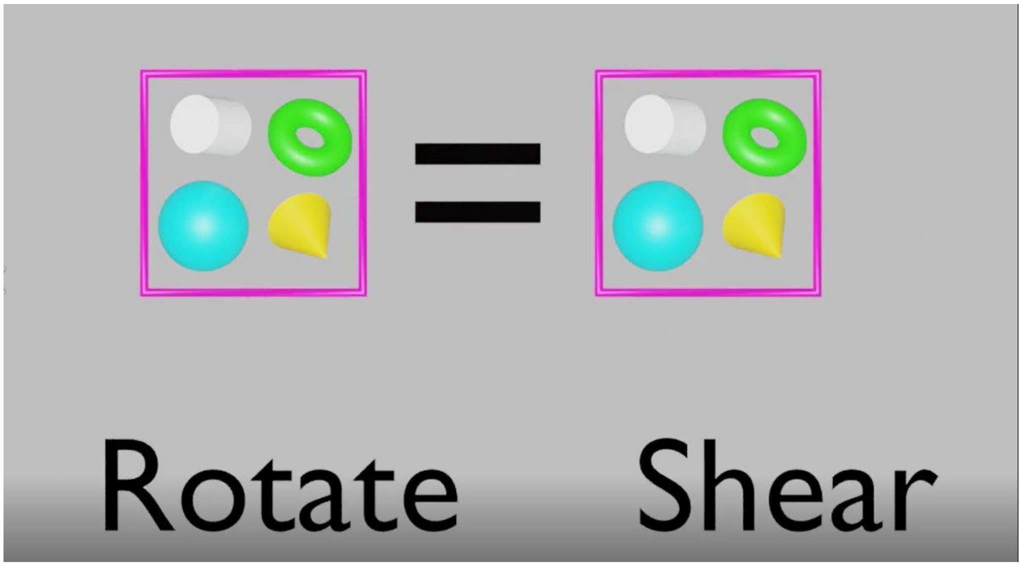


**Supplementary Movie 1** - Animation to illustrate the projection of a rotated and a sheared volume, and illustration of the working principle of the scanning unit.

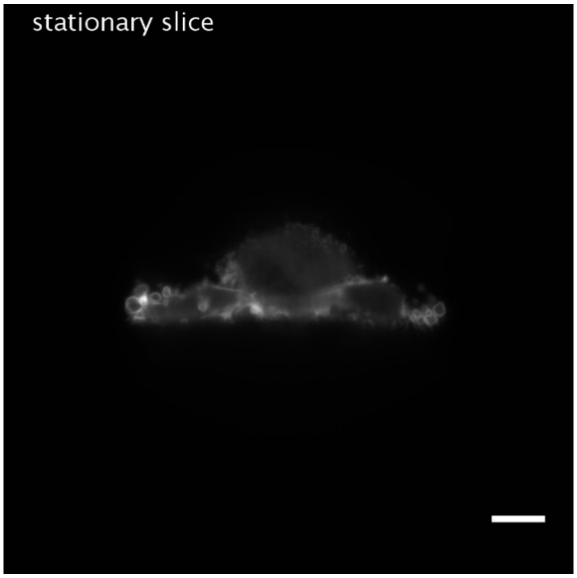


**Supplementary Movie 2 -** Synthesis of projections by increasing scan frequency in a Field Synthesis microscope (sample scanning), without and with lateral shearing. The cell imaged is an MV3 cell expressing AKT-PH GFP. Scale bar: 10 microns

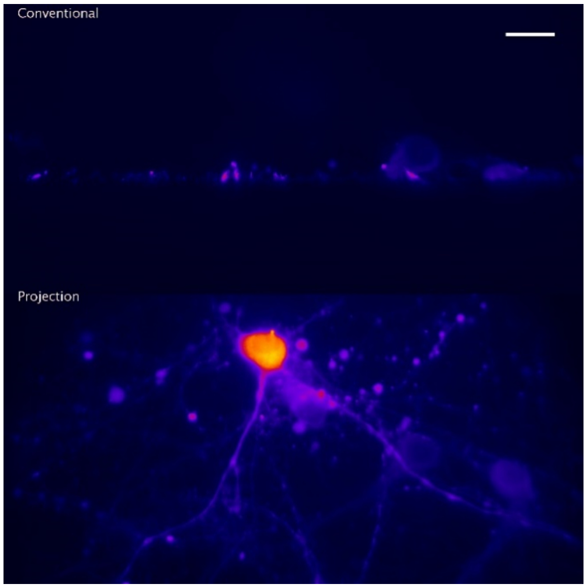


**Supplementary Movie 3 -** Sample exploration in a conventional view in an oblique plane microscope (top), and projection view using OPM-Pro (bottom). Cultured Neurons labeled with GCaMP6F. Scale bar: 20 microns

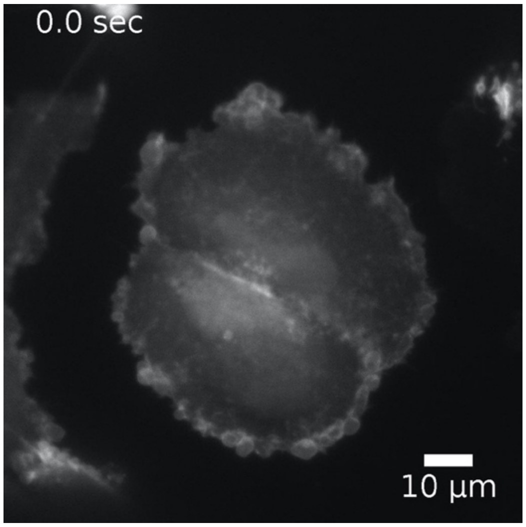


**Supplementary Movie 4 -** Variation of scan and shearing amplitude in projection imaging. MV3 cell expressing AKT-PH GFP as imaged by LLSM-Pro.

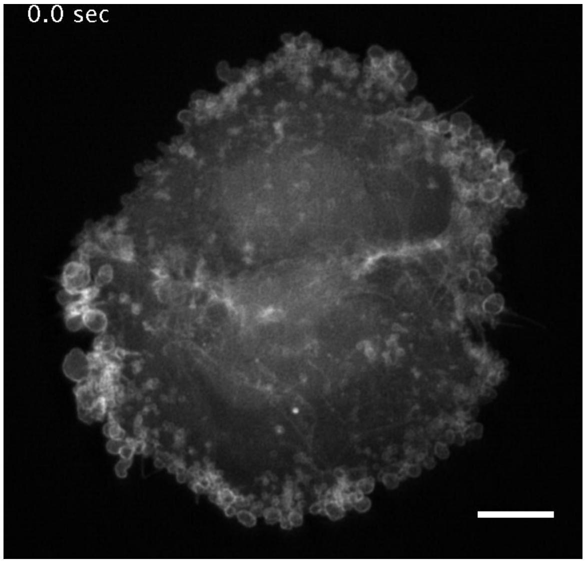


**Supplementary Movie 5 -** Bleb dynamics on an MV3 cell expressing AKT-PH GFP as imaged by LLSM-Pro. Scale bar 10 microns.

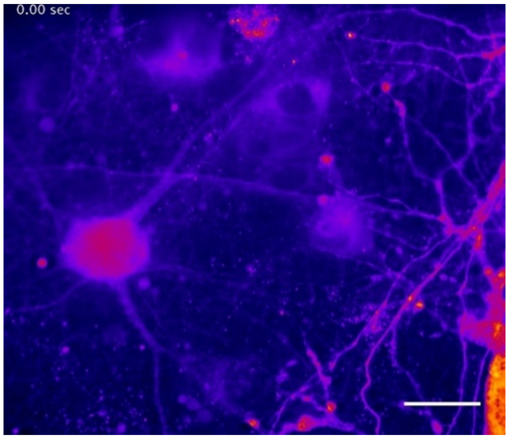


**Supplementary Movie 6 -** Calcium dynamics in cultured Neurons expressing GCAMP-6F, imaged at 12.5Hz Volumetric rate with OPM-Pro. Scale bar: 20 microns

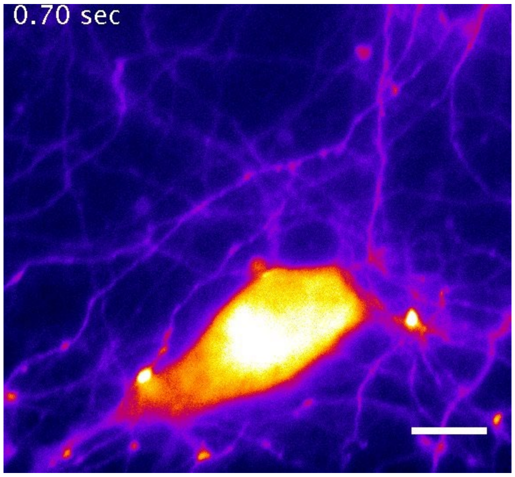


**Supplementary Movie 7 -** Stimulation of cultured neurons expressing GCaMP6F with 100 mM KCI. Imaged at 20Hz volumetric rate with OPM-Pro. Scale bar: 10 microns.

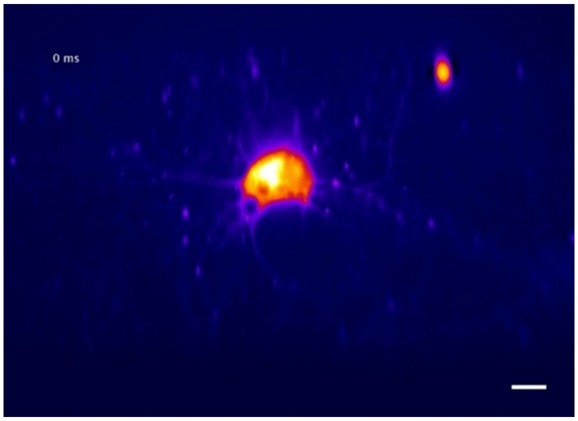


**Supplementary Movie 8 –** Projection imaging of cultured neurons expressing GCaMP6F while varying the viewing angle. 20Hz volumetric imaging rate. Scale bar: 10 microns.

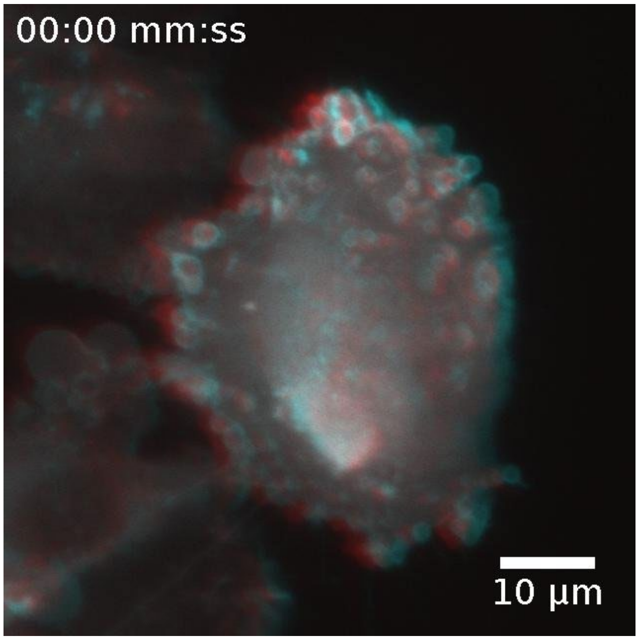


**Supplementary Movie 9 -** Anaglyph for 3D viewing of a MV3 cancer cells expressing AKT-PH GFP. 3D red cyan glasses are recommended to view this image correctly. The movie was created by two different views (15° and -15°) of the cell at a frame rate of 1Hz.

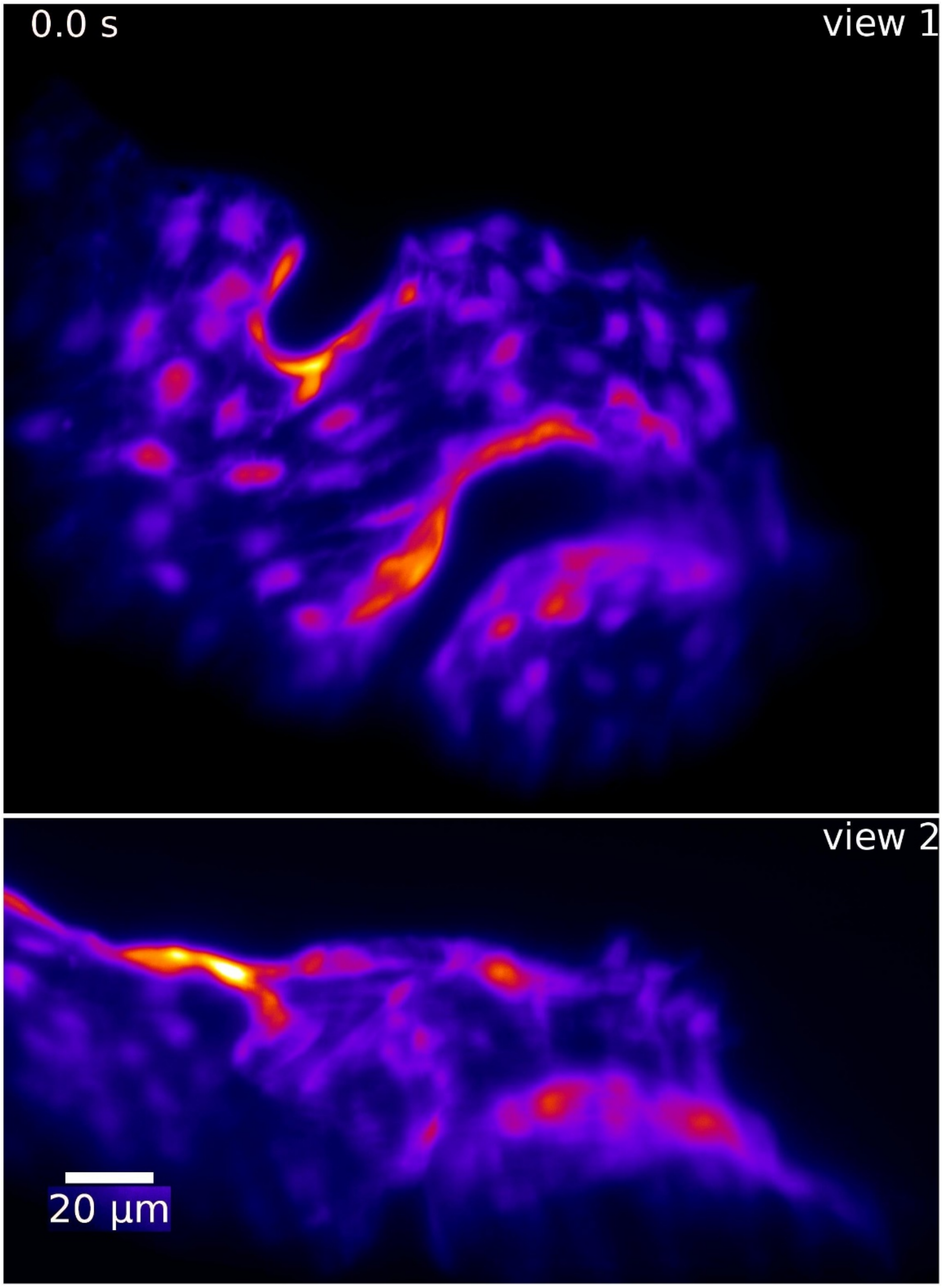


**Supplementary Movie 10 -** Simultaneous dual view imaging of an embryonic zebrafish heart at a framerate of 10Hz. The veins/endothelial tissue of the zebrafish was labeled with GFP (Tg(krdl:GFP)).
